# Local sleep during mind-wandering enhances processes of spatial attention allocation

**DOI:** 10.1101/2020.06.05.136374

**Authors:** Christian Wienke, Mandy V. Bartsch, Lena Vogelgesang, Christoph Reichert, Hermann Hinrichs, Hans-Jochen Heinze, Stefan Dürschmid

**Affiliations:** Department of Neurology, Otto-von-Guericke University, Leipziger Str. 44, 39120 Magdeburg, Germany; Forschungscampus STIMULATE, Otto-von-Guericke University, Universitätsplatz 2, 39106 Magdeburg; Department of Behavioral Neurology, Leibniz Institute for Neurobiology, Brenneckestr. 6, 39120 Magdeburg, Germany; CBBS - center of behavioral brain sciences, Otto-von-Guericke University, Universitätsplatz 2, 39106 Magdeburg, Germany; German Center for Neurodegenerative Diseases (DZNE), Leipziger Str. 44, 39120 Magdeburg

**Keywords:** high frequency activity, local sleep, mind wandering, N2pc, visual spatial attention

## Abstract

Mind wandering (MW) is a subjective, cognitive phenomenon, in which thoughts move away from the task towards an internal train of thoughts, possibly during phases of neuronal sleep-like activity (local sleep, LS). MW decreases cortical processing of external stimuli and is assumed to decouple attention from the external world. Here, we directly tested how indicators of LS, cortical processing and attentional selection change in a pop-out visual search task during phases of MW. Participants brain activity was recorded using magnetoencephalography, MW was assessed via self-report using randomly interspersed probes. As expected, MW worsened performance being accompanied by a decrease in high frequency activity (HFA, 80-150Hz) and an increase in slow wave activity (SWA, 1-6Hz), consistent with the occurrence of LS. In contrast, visual attentional selection as indexed by the N2pc component was enhanced during MW with the N2pc amplitude being directly linked to participants’ performance. This observation clearly contradicts accounts of attentional decoupling predicting a decrease in attention-related responses to external stimuli during MW. Together our results suggest that MW occurs during phases of LS with processes of attentional target selection being upregulated, potentially to compensate for the mental distraction during MW.

## Introduction

Depending on the time spent awake and the richness of experiences rodents and humans enter local sleep-like states, which manifests both as high amplitude slow wave activity (SWA) in the delta/theta range (1-6Hz) and brief neuronal silencing (Vyazovskiy et al. 2011). Phenomenologically local sleep (LS) is assumed to unearth mind-wandering (MW) (Andrillon et al. 2019), during which attention shifts inwards to self-centered matters (Smallwood and Schooler 2006). Both LS and MW increase behavioral errors (Carriere et al. 2008; Smallwood et al. 2008; Bernardi et al. 2015; Seli 2016; Leszczynski et al. 2017) promoting the prediction of perceptual and attentional decoupling (Schad et al. 2012; Christoff et al. 2016). The former is attested by reduced electrophysiological responses (Smallwood et al. 2008; Kam et al. 2011, 2018; Christoff et al. 2016), evidence for attentional decoupling from the environment, however, is limited (Schad et al. 2012). Importantly, since off periods (LS and MW) during waking are potentially harmful (He et al. 2011; Kucyi et al. 2013; Yanko and Spalek 2014; Brandmeyer and Delorme 2018) the survival in general would be endangered if the brain’s need for rest is met entirely during waking (Vyazovskiy and Harris 2013) at the expense of the ability to flexibly shift attention to key features in the environment. Still, how the brain’s ability to shift attention varies during off periods (LS and MW) is unknown.

An established electrophysiological response attributed to the focusing of visual attention onto a target searched among distractors, the EEG component N2pc (Luck and Hillyard 1994a; Eimer 1996; Luck et al. 1997; Hopf et al. 2000; Mazza et al. 2009; Boehler et al. 2011), permits to test this variation. The N2pc is characterized by a more negative deflection at posterior EEG channels contralateral to the visual field in which the target was presented. Theoretically there are at least two principal scenarios which can be tested using the N2pc. On the one hand, the attentional decoupling account predicts that the N2pc as an index of attentional selection gradually decreases with MW. On the other hand, it could be hypothesized that the N2pc increases with MW since MW and external distractors are assumed to share a common underlying mechanism (Forster and Lavie 2014; Unsworth and McMillan 2014) and the N2pc increases with an increasing amount of distracting information (Mazza et al. 2009).

Using the high spatiotemporal and spectral resolution of magnetoencephalographic recordings (MEG) we investigated how cortical dynamics varied with self-reports ranging from being ON (uninterrupted focus on the external environment) to OFF (MW) the task, in which subjects searched for a color-defined pop-out (target) among task-irrelevant distractors. Moreover, we hypothesized that if associated with LS, MW leads to SWA and neuronal silencing. The latter we would expect to be reflected in a reduction in high frequency activity (HFA, 80-150Hz), a correlate of population neural firing rate (Mukamel et al. 2005; Liu and Newsome 2006; Manning et al. 2009; Miller et al. 2009; Ray and Maunsell 2011) and preferred proxy for asynchronous areal activation (Miller et al. 2009, 2014; Privman et al. 2013; Coon and Schalk 2016; Kupers et al. 2017) ideally suited to test neuronal silencing.

## Materials and Methods

### Participants

Sixteen subjects (5 female, range: 18-39 years, *M*: 27.13, *SD*: 5.85) participated after providing their written informed consent. One subject who did not experience MW was excluded. All participants reported normal or corrected-to-normal vision and none reported any history of neurological or psychiatric disease. All recordings took place at the Otto-von-Guericke University of Magdeburg and were approved by the local ethics committee (“Ethical Commitee of the Otto-von-Guericke University Magdeburg”) and each participant was compensated with money.

### Paradigm

Participants were presented with a stimulus array of red, green, and blue grating patterns each consisting of 3 colored and 2 grey stripes viewed through a circular aperture (Fig ***1***). The grey stripes matched the grey of the background. While either of the green and red gratings served as target, blue gratings always served as distractor items. Stimulus arrays consisted of 18 gratings arranged in two blocks of 9 gratings left and right below the fixation cross. Presentation of search displays in the lower visual field has been shown to evoke a stronger N2pc amplitude (Luck et al. 1997; Hilimire et al. 2011). Participants were instructed to keep fixation on the fixation cross located at 1.9° visual angle (va) above the stimulus array. The size of each grating was 1.15° va, distance between single gratings (edge-to-edge) was 0.69° va. The left and right block of gratings each had a size of 4.83° by 4.83° va, the horizontal distance between both blocks (inner edges) amounted to 5.15° va. Diagonal distance between the fixation cross and the center of the nearest upper grating was 2.81° va. Target gratings could be tilted left or right in ten steps of 1.5°, with the smallest tilt being 1.5° and the maximal tilt being 15° from the vertical axis. Orientation and tilt angle of the non-target and distracter gratings varied randomly. Stimulus generation and experimental control was done using Matlab R2009a (MATLAB and Statistics Toolbox Release 2009, The MathWorks, Inc., Natick, Massachusetts, United States.) and the Psychophysics Toolbox (Brainard 1997; Pelli 1997; Kleiner et al. 2007). Colors were matched for isoluminance using heterochromatic flicker photometry (Lee et al. 1988).

**Figure 1.**
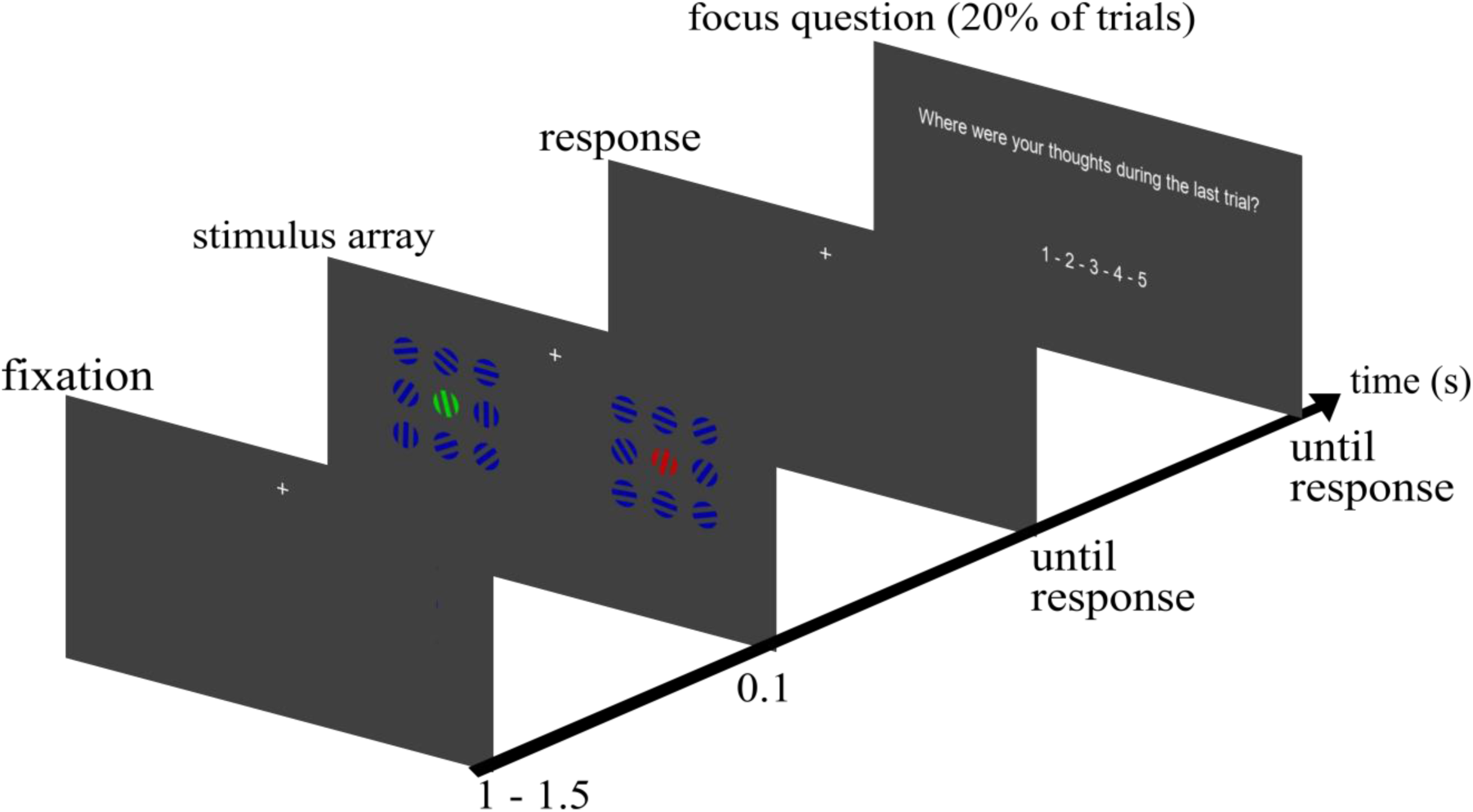
Single trial with focus question (see text for detail)

### Procedure

At the beginning of each of the 12 blocks, participants were instructed to attend either only to the red or green grating and report via button press towards which side it was tilted (left: index finger, right: middle finger of the right hand). Target color assignment alternated blockwise. In blocks with the red grating as target the green grating served as non-target and had to be ignored and vice versa. The target could appear at each of the eighteen locations. The location of the non-target was constrained to the mirrored location in the opposite grating block to keep equal distances to the fixation cross for both target and non-target gratings. Each trial started with a fixation period of 1250 msec (±250msec) before the stimulus array was presented for 100 msec. Participants were asked to respond as fast and accurately as possible. Afterwards the next trial started. The experiment started with a training block of twenty trials to familiarize participants with the procedure. After twenty consecutive trials, a blinking pause allowed participants to blink and rest their eyes. These pauses lasted seven seconds. Each block consisted of 100 trials.

### Experience sampling

Throughout the experiment we delivered thought probes in pseudorandomly chosen trials (20%) asking participants to rate their attentional focus, in the period immediately prior to the probe, on a five point scale from 1 (“thoughts were anywhere else” – OFF) to 5 (“thoughts were totally at the task” – ON). Responses to focus questions were given with all five fingers of the left hand (thumb: 5, index finger: 4, middle finger: 3, ring finger: 2, little finger: 1). The probes were presented following orientation discrimination, with the restriction that two probes were separated by a minimum of one intervening search trial. The probes were initiated by an auditory stimulus (500 Hz, ca. 85 dB for 200 msec). To increase statistical power we grouped the five MW ratings in three groups of mental state (OFF: 1&2, MID: 3, ON: 4&5). Statistical analyses between mental states were performed on this subset of trials.

### MEG recording

Participants were equipped with metal-free clothing and seated in a dimmed, magnetically shielded recording booth. Stimuli were presented via rear projection onto a semi-transparent screen placed at a viewing distance of 100cm in front of the participants with an LCD projector (DLA-G150CLE, JVC, Yokohama, Japan) that was positioned outside the booth. Responses were given with the left and right hand via an MEG compatible LUMItouch response system (Photon Control Inc., Burnaby, DC, Canada). Acquisition of MEG data was performed in a sitting position using a whole-head Elekta Neuromag TRIUX MEG system (Elekta Oy, Helsinki, Finland), containing 102 magnetometers and 204 planar gradiometers. Sampling rate was set to 2000Hz. Vertical EOG was recorded using one surface electrode above and one below the right eye. For horizontal EOG, one electrode on the left and right outer canthus was used. Preparation and measurement took about 2 hours.

### Preprocessing and artifact rejection

We used MatLab 2013b (Mathworks, Natick, USA) for all offline data processing. The 102 magnetometers were involved in our analyses. All filtering (see below) was done using zero phase-shift IIR filters (4^th^ order; filtfilt.m in Matlab). First, we filtered the data between 1 and 200 Hz and used a threshold of 3pT, which the absolute MEG values must not exceed, to discard trials (−1 sec to 2 sec around stimulus onset – sufficiently long to prevent any edge effects during filtering) of excessive, non-physiological amplitude. We then visually inspected all data, excluded epochs exhibiting excessive muscle activity, as well as time intervals containing artifactual signal distortions, such as signal steps or pulses. We refrained from applying artifact reduction procedures that affect the dimensionality and/or complexity of the data like independent component analysis. Time series of remaining trials were used to characterize HFA (80-150 Hz), SWA (1-6Hz) and the N2pc (1-30Hz, main frequency range for cognitive event-related-potential (ERP) components, see (Luck 2005)). Resulting time series were used to characterize brain dynamics over the time course of visual target detection. Each trial (−1 to 2 sec around stimulus onset) was baseline corrected relative to the 200 msec interval prior to the stimulus onset.

### Statistical analysis

To correct statistical significance for multiple comparisons we compared each statistical parameter against a surrogate distribution, which were constructed by randomly yoking labels of the trials and repeating the ANOVA, t-tests, and Pearson’s correlation coefficient. Consequently, reported p-values represent the statistical significance relatively to the constructed surrogate distribution.

### I Behavioral results

We tested whether the ratio of ON and OFF ratings changed across the experiment to rule out the possibility that changes in cortical dynamic are a result of a change across the experiment and not of fluctuations of the mental state throughout the experiment. We divided the 12 experimental blocks in 4 parts by averaging ratings in 3 consecutive blocks since individual subjects did not make use of each of the five ratings in single blocks and compared the number of ON and OFF ratings across these 4 parts with a 4×2 ANOVA with the factors block (I,II,III, and IV) and mental state (ON vs. OFF).

Performance, measured as percent correct responses, was averaged across tilt angles for each subject and compared between mental states with a one-way ANOVA. Performance during focus trials was then correlated with N2pc (see below) amplitude to test whether N2pc strength predicts performance.

Reaction times (RTs) were grouped for the three mental states and averaged across subjects. The averaged RTs where then compared using a one-way ANOVA with the factor mental state (OFF, MID, ON).

### II HFA response (neuronal silencing)

We then obtained the HFA response. For each trial we band-pass filtered each magnetometer’s time series in the broadband high frequency range (80-150 Hz). We obtained the analytic amplitude *A*_*f*_(*t*) of this band by Hilbert-transforming the filtered time series. In the following, HFA refers to this Hilbert transform. We smoothed the HFA time series such that amplitude value at each time point *t* is the mean of 25 msec around each time point *t*. We then baseline-corrected by subtracting from each data point the mean activity of the 200 msec preceding the stimulus onset in each trial and each channel. We then identified stimulus-responsive channels showing a significant (compared to an empirical distribution, see below) amplitude modulation in the HFA following the onset of the visual search array. We first calculated the average activity modulation *Ā*_*HFA*_ averaged across the 300 msec following the stimulus onset from which we subtracted the baseline activity 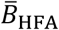 preceding the stimulus onset. The difference between 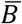 and *Ā* was compared against a surrogate distribution. In each iteration, time series of each channel were circularly shifted between −500 msec and 300 msec separately, and new (surrogate) trial averages (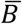 and *Ā*) were calculated. Channels exceeding the 97.5^th^ percentile of the channel specific surrogate 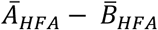 distribution were classified as showing a significant HFA modulation following stimulus onset. Second, to test for HFA differences between mental states, a one-way ANOVA (OFF, MID, ON) was conducted at each time point between 100 msec pre- and 500 msec post-stimulus. The F-value of the main effect “mental state” parameterizes neuronal silencing in the HFA response, with high F-values indicating a large difference in HFA amplitude between mental states. To set a threshold for significant difference, an empirical distribution of the main effect was constructed by randomly reassigning the labels (OFF – MID – ON) to the single trials in 1000 permutations. Peak responses (maximal average HFA response following stimulus onset) in each of the mental states were compared against a surrogate distribution. In each iteration, time series of each channel were circularly shifted time series of participants between −500 msec and 300 msec separately, and new (surrogate) trial averages were calculated. From these trial averages we calculated the peak value in the time range of 0 to 300 msec following stimulus onset. Mental states exceeding the 97.5^th^ percentile were classified as showing significant HFA modulation.

### III High amplitude slow wave oscillation

For each trial we band-pass filtered each magnetometer’s time series in the frequency range of slow wave oscillations (1-6 Hz) and z-scored the obtained analytic amplitude *A*_*f*_(*t*) of this band by Hilbert-transforming the filtered time series. In the following, SWA refers to this Hilbert transform. We then counted the number of peaks of the SWA defined as local maxima exceeding 3 SD in each trial at each channel. Next, we identified channels with a high number of SWA peaks. To this end we compared the average number of SWA peaks across subjects against a surrogate distribution. In each of 1,000 iterations we randomly exchanged channel labels in each subject and new (surrogate) channel averages were calculated across participants. Channels exceeding the 97.5^th^ percentile of the channel specific surrogate distribution were classified as showing a significant SWA modulation following stimulus onset (SWA channels). The number of SWA peaks were averaged separately for the three mental states across SWA channels in each participant. We then carried out a one-way ANOVA with factor mental state (OFF – MID – ON) at each time point, with single participants as random variable. The F-value of the main effect “mental state” parameterizes the occurrence of SWA with high F-values indicating a large difference in the number of SWAs between mental states. To set a threshold for significant difference, an empirical distribution of the main effect was constructed by randomly reassigning the labels (OFF – MID – ON) to the single trials in 1000 permutations.

### IV N2pc

The N2pc was calculated from the subset of trials in which a focus question was presented. First, using t-tests, we compared for each subject the magnetic response (1-30Hz) at each sensor for targets in the left visual field (LVF) vs targets in the right visual field (RVF) at each time point, irrespective of target color and distance to fixation cross. Subtracting the RVF response from the LVF response, as done by the t-test, removes activity that is solely based on sensory processes since all trials contain a red and a green pop-out grating (Hopf et al. 2000). From these distributions of t-values, occipito-temporal sensors showing maximal positive and maximal negative t-values in the time range from 200 msec to 300 msec post-stimulus were then selected individually for each subject on each hemisphere. These two channels on each hemisphere were then combined by subtracting the response of the influx-channel from the efflux-channel (Max_positive_ - Max_negative_) separately for targets in the LVF and RVF. The N2pc for each hemisphere was finally extracted from this combined signal by subtracting the average for targets in the RVF from the average for targets in the LVF. Using the individually selected sensors we then extracted the N2pc for the three mental states accordingly.

To rule out hemispherical differences in N2pc amplitude, we conducted a t-tests at every time point between the N2pc elicited over left and right hemisphere. Results were compared against a distribution derived from randomly reassigning the sides and repeating the t-test in 1000 iterations. To anticipate, our time resolved t-test did not reveal differences between hemispheres hence we collapsed N2pc responses across hemispheres. In the next step we tested whether the N2pc was significantly elevated over baseline. We baseline-corrected the N2pc time series of each subject by subtracting from each data point the mean activity of the 200 msec preceding the stimulus onset. We then tested whether the average N2pc shows a significant (compared to an empirical distribution, see below) amplitude modulation following the onset of the visual search array. We first calculated the average activity modulation *Ā*_N2pc_ averaged across the 200-300 msec following the stimulus onset from which we subtracted the baseline activity 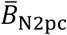 preceding the stimulus onset. The difference between 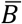 and *Ā* was compared against a surrogate distribution. In each iteration, time series of each subject were circularly shifted between −500 msec and 300 msec separately, and new (surrogate) trial averages (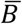 and *Ā*) were calculated. Time points exceeding the 97.5^th^ percentile of the channel specific surrogate 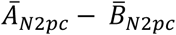 distribution were classified as showing a significant N2pc modulation following stimulus onset. The first time point of significant N2pc modulation in each subject was used as N2pc onset. Using a time point – by – time point ANOVA between −100 and 600 msec with the factor mental state (OFF, MID, ON) we tested whether the N2pc differs between focus conditions. The F-value of the main effect “mental state” parameterizes the variation of the N2pc as a function of mental states with high F-values indicating a large difference in N2pc amplitude between mental states. To set a threshold for significant difference, an empirical distribution of the main effect was constructed by randomly reassigning the labels (OFF – MID – ON) to the single trials in 1000 permutations.

### VI Local sleep-N2pc correlation

First, HFA and N2pc onset times were compared via t-test to analyze temporal discrimination between both. Second, to examine the interaction between HFA and N2pc over the different mental states, HFA and N2pc time series were averaged separately for the three mental states in each participant for the interval between onset and offset (interval between significant elevation over baseline). We then carried out a two-way ANOVA with factor MEG response (N2pc – HFA) and mental state (OFF – MID – ON) at each time point, with single participants as random variable. Third, for each mental state N2pc (averaged across the interval of significant amplitude modulation for all trials) was correlated with HFA response (averaged across the interval of significant amplitude modulation for all trials). The resulting Pearson’s correlation values were tested against a surrogate distribution. This surrogate distribution was constructed by randomly assigning the HFA values of each participant with the N2pc values from another participant in 1000 iterations.

## Results

### I Behavioral results

MW ratings differed in frequency (*F*_2,42_ = 10.11, *p* < 0.001; ON 51.25% (*SD:* 27%), MID 33.1% (*SD:* 18.7%), and OFF 15.67% (*SD:* 16.8%); ***Fig. 2A***) with more ON than MID ratings (*t*_*14*_ = 2.21, *p* = .035) and more MID than OFF ratings (*t*_*14*_ = 2.56, *p* = 0.016). The ratio of ratings did not vary across blocks: main effect of block (*F*_3,112_ = 0.03, *p* = .99) and interaction (*F*_3,112_ = 0.6; *p* = .6) were not significant (***Fig. 2A***). While ON ratings did not vary across blocks (all *p*’s > .1), OFF ratings increased from block I to II (*t*_*14*_ = 2.5; *p* = .02) but remained constant afterwards. Performance varied with mental state (*F*_2,42_ = 5.14, *p* = .01) with worse performance during OFF trials (*M*: 70.2%, *SD*: 18.8%) than during MID trials (*M*: 80.2%, *SD*: 7%; *t*_14_ = 2.62, *p* = .01) or ON trials (*M*: 84.7%, *SD*: 7%; *t*_14_ = 2.09, *p* = .03). No differences were observed between ON and MID trials (*t*_14_ = 1.76, *p* = .1; ***Fig. 2B***). Also, reaction times differed significantly between mental states (*F*_2,42_ = 2.75 *p* = 0.0031) with slower RTs during OFF (*M*: 898 msec, *SD*: 1028 msec) compared with ON (*M*: 433 msec, *SD*: 146 msec; *t*_*14*_ = 1.72, *p* = .04), a trend of statistical significance between OFF and MID trials (*M*: 489 msec, *SD* 212 msec; *t*_28_ = 1.48, *p* = .07), but no differences between ON and MID trials (*t*_28_ = 0.87, *p* = .38; ***Fig. 2D***).

**Figure 2.**
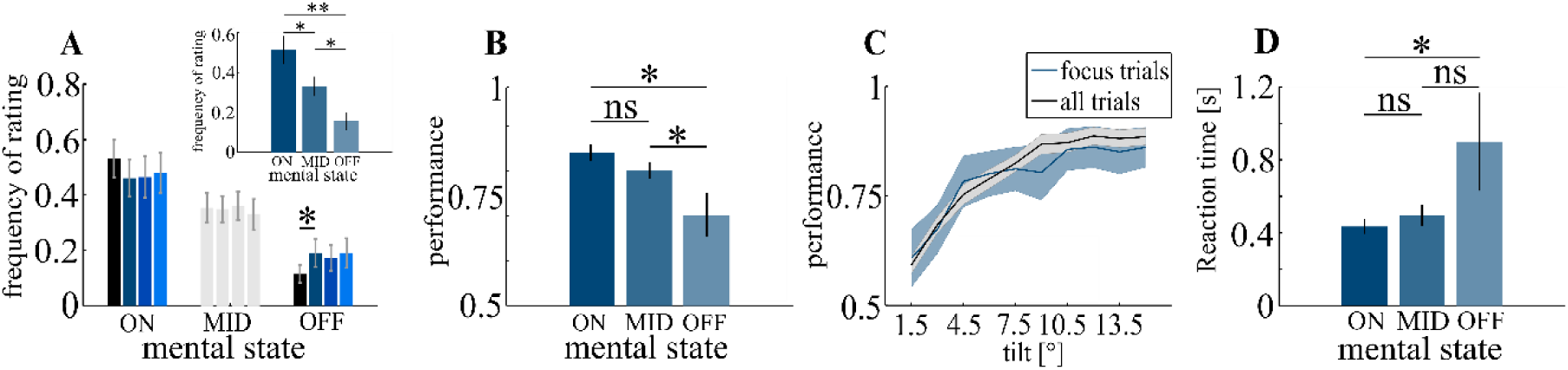
Behavioral data, **A:** participants made more ON and MID than OFF ratings (small inset). Only between the first and the second quarter of the experiment was a significant increase in OFF ratings, which then remained constant. **B:** subjects made more errors during OFF trials than during ON and MID trials. **C:** performance varied between tilt angles across all trials (black) and across the subset of trials after which a focus question was presented (blue). **D:** Reaction times were significantly longer in OFF vs. ON trials. Errorbars and shaded areas represent the standard error of the mean (SEM).* p < 0.05, ** p < 0.01

### II HFA response (neuronal silencing)

15 occipital magnetometers showed stimulus response in the HFA between 81 and 234 msec post-stimulus (HFA_max_ = 1.24fT at 161 msec, *p* < .001, ***Fig. 3A***,***B***,***C***). The HFA differed between mental states between 145 and 171 msec post-stimulus (F_crit_ = 2.74; *F*_max_ = 3.18 at 151msec, *p* = .02, ***Fig. 3D***) with smaller HFA in OFF (*M*: .47fT, *SD*: .93fT) vs. ON (M: 1.24fT, SD: .82fT; *t*_*14*_ = 2.16, *p* = .02) and vs MID trials (*M*: 1.25fT, *SD*: 1.28fT; *t*_*14*_ = 2.04, *p* = .03) but no difference between ON and MID (*t*_*14*_ = 0.53, *p* = .69). Importantly, in contrast to ON (critical peak amplitude = .63fT, HFA_max_ = 1.29fT at 149 msec; p < .001) and MID trials (HFA_max_ = 1.33fT at 152 msec; p < .001), HFA did not show significant peak response in OFF trials indicating that HFA completely vanished (HFA_max_ = .5fT at 151 msec, *p* = .15).

**Figure 3:**
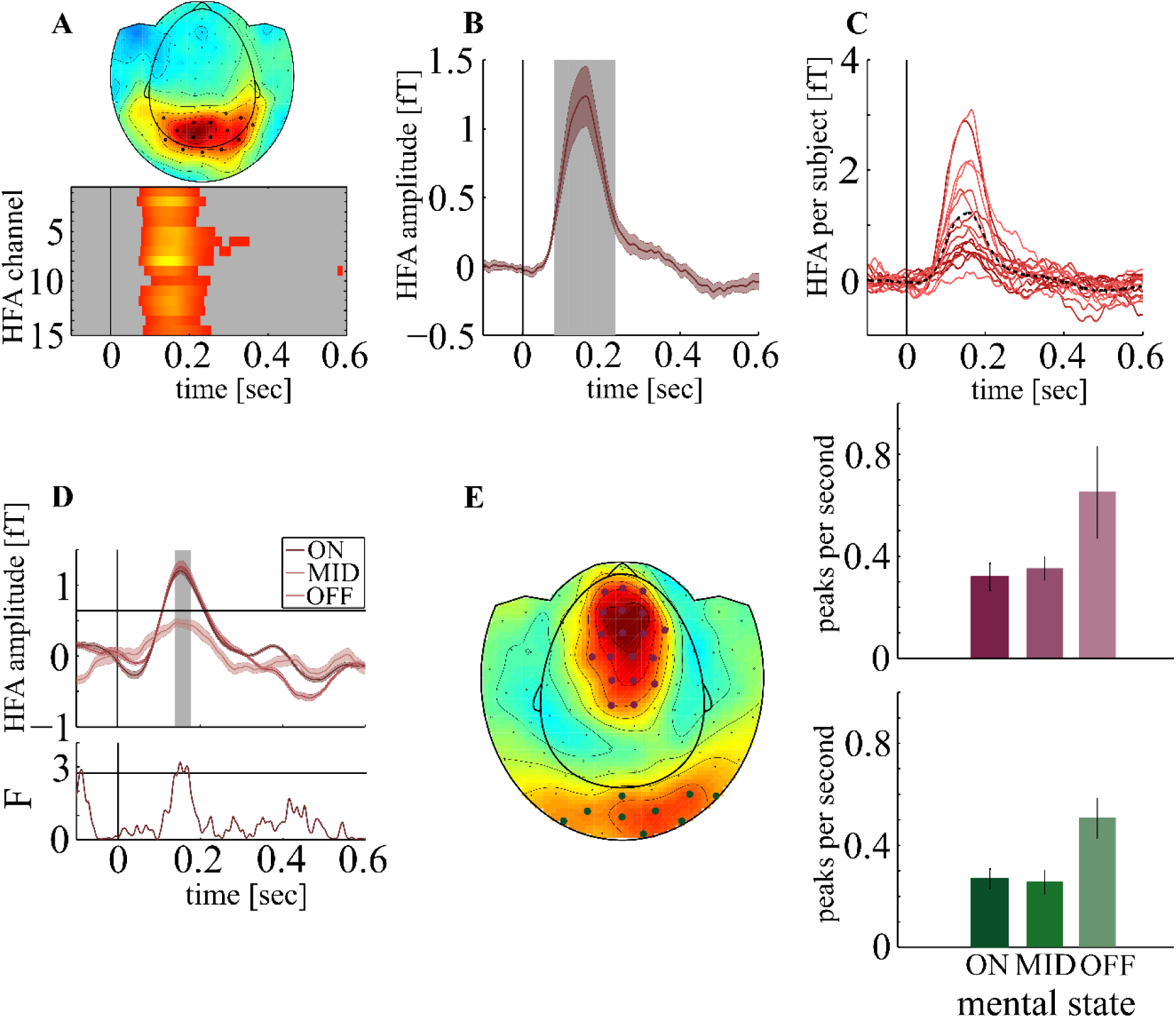
HFA **A:** Grand Average ERMF (80-150Hz) averaged across all focus trials and subjects between 100 and 200 msec post-stimulus (top) shows 15 occipital sensors with significant response after stimulus onset. HFA onset and time course (bottom) are highly similar. **B:** Averaged across all trials and subjects, we found a HFA between 81 and 234 msec post-stimulus (gray inset). **C:** HFA response averaged across significant sensors for each subject. Dotted black line represents average across subjects. **D top:** HFA for each mental state, averaged across subjects. Gray inset represents time of significant differences in amplitude between mental states. Horizontal line represents critical peak amplitude modulation. **D Bottom:** Time course of F-values. Horizontal line represents critical F-value for statistical significance. **E:** 28 Sensors showed significant SWA (left). The Number of SWA peaks in occipital sensors (green, lower right) was significantly elevated during OFF trials (red: frontal sensors). Vertical lines represent stimulus onset. Shaded Areas around curves represent SEM.

### III High amplitude slow wave oscillations

28 MEG sensors covering a frontal-parietal (N_crit_ = .3Hz; N_SWA_ = .43Hz; *p* < .0001) and an occipital channel cluster (N_SWA_ = .38; *p* < .0012, ***Fig. 3E***) showed a significant number of SWA. In frontal-parietal sensors we observed a trend towards differences in frequency of SWA between mental states (*F*_2,42_ = 2.7; *p* = .07, Fig ***3E***), but a highly significant difference in occipital sensors (*F*_2,42_ = 5.9; *p* < .0001, Fig ***3E***) with more SWA peaks in OFF (N_SWA_ = .51) vs ON (N_SWA_ = .27; *t*_*14*_ = 3.4; p = .004) and vs MID trials (N_SWA_ = .25; *t*_*14*_ = 2.6; p = .02) in the occipital region.

### IV N2pc

Attentional target selection elicited an N2pc between 179 and 319 msec post-stimulus (N2pc_crit_ = 4fT, N2pc_max_ = 61.7fT at 258 msec, *p* < .001; ***Fig. 4A***,***B***) with no differences between hemispheres (*t*_crit_ = ±2.74, *t*_max_ = −1.74 at 71 msec, *p* = .94). The N2pc differed between mental states between 213 and 298 msec post stimulus (*F*_crit_ = 3.53, *F*_max_ = 7.62 at 256 msec post-stimulus, *p* < .001; ***Fig 4C***,) with a larger amplitude in OFF (*M:* 78.69fT, *SD:* 46.16) vs MID (*M:* 50.65fT, *SD:* 28.89; *t*_*14*_ = 3.44, *p* = 0.01) and vs ON (*M:* 38.82fT, *SD:* 19.73; *t*_*14*_ = 4.1, *p* = .002) but no significant difference between ON and MID trials (*t*_*14*_ = 0.39, *p* = .69).

**Figure 4:**
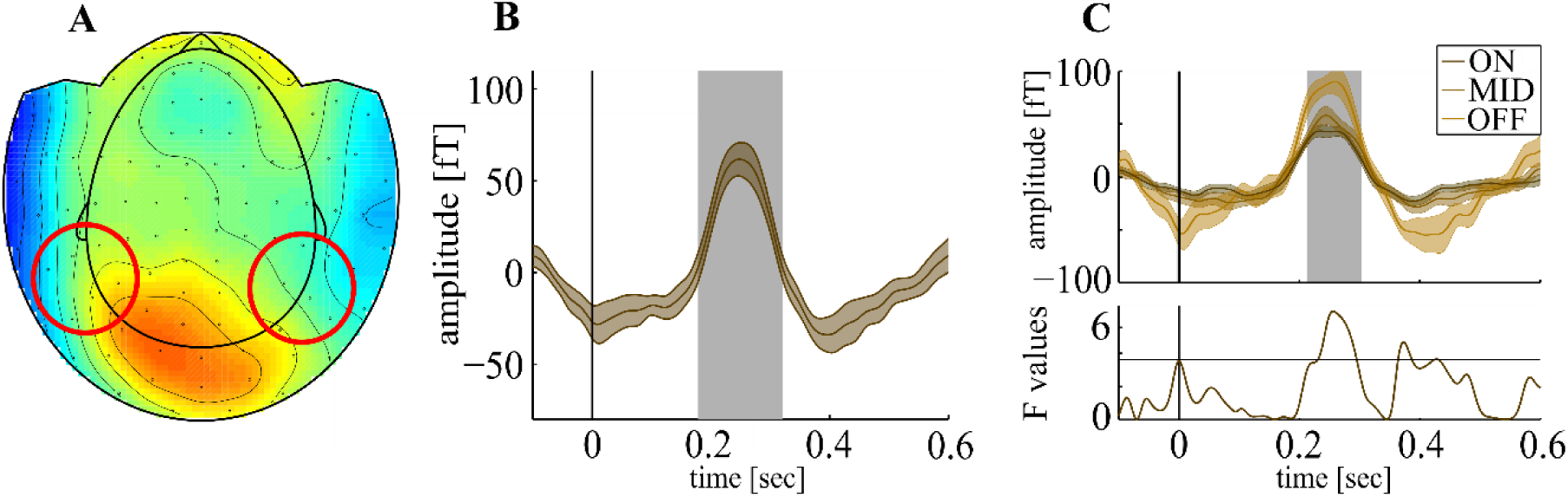
N2pc **A:** Grand average event related magnetic field (ERMF; 1-30Hz) averaged across analyzed trials between 200 and 300 msec post-stimulus. Circles represent probable location of underlying dipoles. **B:** N2pc averaged across analyzed trials and subjects. We found a significant N2pc between 179 and 319 msec post-stimulus (gray inset). **C top:** N2pc for each mental state, averaged across subjects. We found significant differences in N2pc amplitude between mental states (gray inset) between 213 and 298 msec post-stimulus. **C Bottom:** time course of F-values. Horizontal line represents critical F-value. Vertical lines represent stimulus onset. Shaded areas around curves represent SEM.

### V Local sleep-N2pc correlation

The number of SWA peaks correlated with the N2pc in OFF trials both in the fronto-parietal and the occipital channel cluster (*r*_*crit*_ = .53, fronto-parietal: *r* = .71; *p* = .0044; occipital: *r* = .6; *p* = .014) but not in ON or MID trials (*r* values range between −.04 to .45; *p* > .025, ***Fig 5A***).

**Figure 5:**
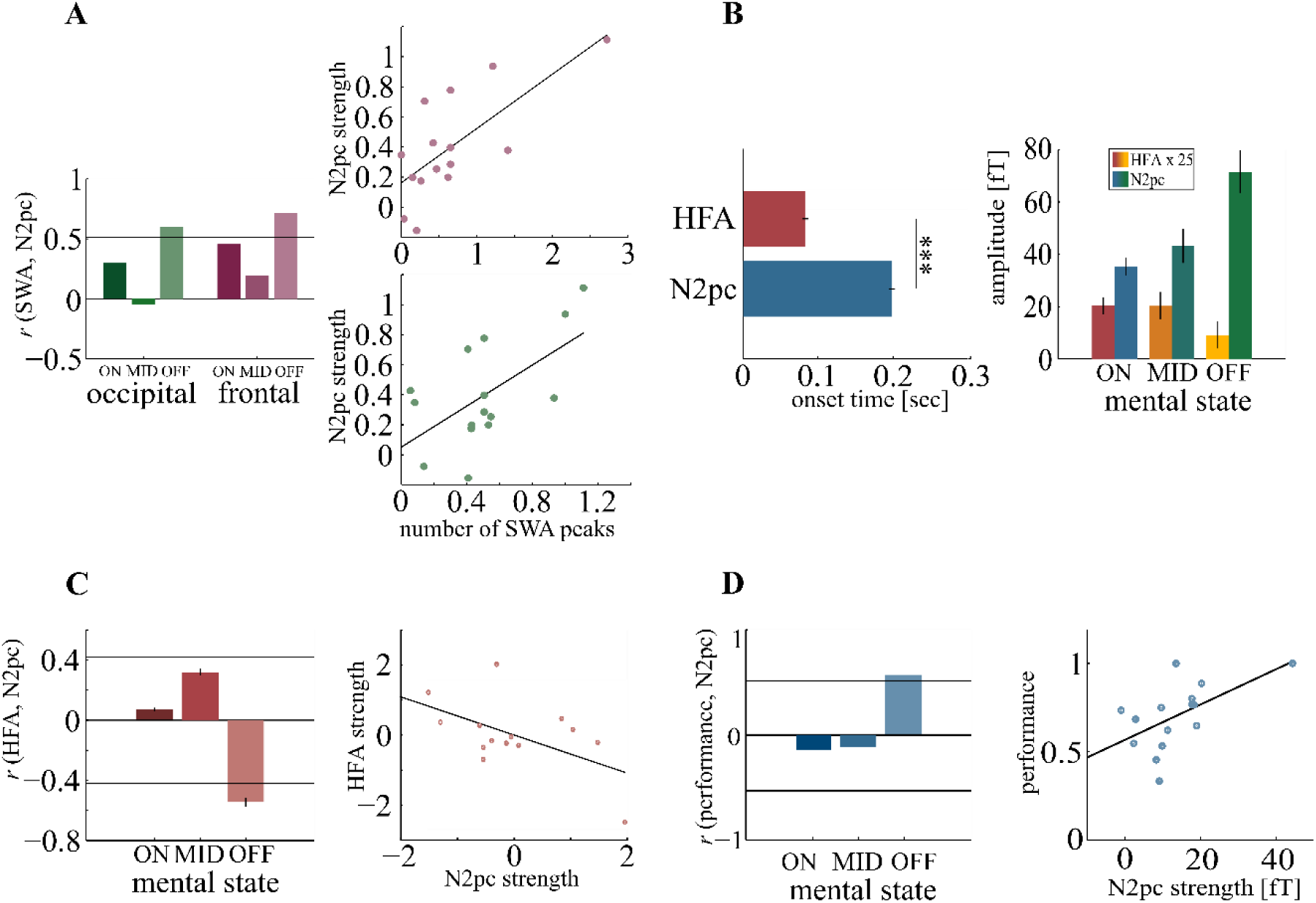
Local sleep-N2pc correlation **A**: Correlation between SWA count and N2pc amplitude was significant only during OFF trials. Horizontal line represents critical correlation value (left). Scatterplots showing the correlation between SWA count and N2pc for OFF trials in frontal (red, upper) and occipital sensors (green, lower)(right). **B:** Onset times for HFA and N2pc differed significantly (left). Average HFA and N2pc amplitude for each mental state. Note that the HFA is scaled up in this plot to compensate for lower amplitudes (right). **C:** Correlation between HFA and N2pc reached significance only during OFF trials. Horizontal lines represent critical correlation values (left). Scatterplot showing the correlation between HFA and N2pc during OFF trials (right). **D:** Correlation between performance and N2pc reached statistical significance only during OFF trials. Horizontal lines represent critical correlation values (left). Scatterplot showing the correlation between performance and N2pc strength during OFF trials (right). Errorbars represent the SEM. *** p < .001

Importantly, the HFA (reflecting initial visual response) showed a significantly earlier onset than the N2pc (HFA: 83 msec post-stimulus, SD: 14 msec; N2pc: 198 msec post-stimulus, SD: 17 msec; *t*_*14*_ = 20.1, *p* < .001, Fig ***5B, left***). Average HFA and N2pc showed a strong interaction with mental states with the N2pc increasing with decreasing HFA (*F*_2,87_ = 11.17, *p* < .001; ***Fig 5B, right***). Similarly to SWA, only in OFF trials HFA correlated with the N2pc (*r*_crit_ = ±.42, *r* = −.54, *p* = .04), indicating that a low HFA amplitude is associated with an increased N2pc amplitude but not in ON (*r* = .07, *p* = .71) or MID trials (*r* = .31, *p* = .27, Fig ***5C***). This enhancement of the N2pc appeared to be behaviorally relevant as in OFF trials, the N2pc was correlated to performance (*r*_crit_ = ±.53, *r* = .57, *p* = .02) but not in ON (*r* = −.14, *p* = .29) or MID trials (*r* = −11, *p* = .33; Fig ***5D***).

## Discussion

We examined the role of local sleep (operationalized as HFA reduction and SWA increase) in the generation of MW, and its impact on spatial attentional allocation. Participants performed a visual search paradigm, yielding robust increases in the HFA response in occipital MEG sensors, followed by the N2pc responses reflecting target selection. When subjects subjectively experienced MW, the HFA response vanished corroborating neuronal silencing (Vyazovskiy et al. 2011) and establishing a direct link between local sleep and MW. In parallel, the number of SWA periods increased with MW, consistent with participants experiencing phases of local sleep. In line with previous studies, performance decreased with manual reaction times being substantially prolonged during MW. In contrast, neural markers of attentional selection were even more pronounced during MW and closely linked to behavioral responses. That is, even though low in performance during OFF trials, subjects showing a higher N2pc amplitude performed better than those with a less pronounced N2pc. In general, during MW and commensurate with signatures of local sleep, processes of attentional target selection, as indexed by the N2pc, rather increased potentially compensating for mental distraction.

Grating stimuli reliably evoked high frequency activity in our non-invasive MEG recordings strongly resembling HFA responses in intracranial recording with a modulation over baseline between 50 and 350 Hz, a fast increasing flank peaking around 200 msec, and a slowly decreasing flank in early visual cortex (Burke et al. 2014; Szczepanski et al. 2014; Golan et al. 2016, 2017; Gerber et al. 2017; Helfrich et al. 2018; Bartoli et al. 2019). The high similarity of the HFA response across subjects indicates that MEG in contrast to EEG can reliably pick up high frequency activity responses to visual stimuli which even has been shown at the single trial level (Westner et al. 2018).

HFA reduction during MW might not result from attentional decoupling but rather reflects neuronal silencing. Previous studies showed reduced electrophysiological responses during MW (Christoff et al. 2016) potentially due to attentional decoupling during MW without deciphering the causal relation between MW and reduced cortical responses. It is assumed that MW attenuates the cortical response (Christoff et al. 2016) – the HFA – since attentional resources are shifted inwards (Smallwood and Schooler 2006) in line with an attentional decoupling account. However, we hypothesize that participants experience MW, since use-dependent neuronal silencing reduces sensory representation of the visual environment in the first place for the following reasons. First, any attentional reduction of the HFA should also predominantly be found in fronto-parietal structures (Szczepanski and Kastner 2013; Szczepanski and Knight 2014; Perrone-Bertolotti et al. 2020) where we did not find any strong stimulus-driven modulation in our study. Second, and most importantly, attentional modulation of cortical responses are amply attested with a reduction of responses (Smallwood et al. 2008; Kam et al. 2011, 2018) often using a contrast between task relevant vs. irrelevant stimuli (Müsch et al. 2014). But task-irrelevant stimuli evoked a comparable HFA response even though smaller in amplitude. Also, in audition even though ignoring the stimulation and attending a second task clear stimulus-driven responses can be seen in frontal and temporal cortex (Dürschmid et al. 2016). Hence, although modulated by attention, ERPs and HFA response in previous studies were preserved. In contrast, we found HFA increase in occipital MEG sensors onsetting as early as ∼90 msec and, most importantly, during MW the HFA vanishes. Hence, HFA reduction is most likely not driven by attention but rather corresponds with neuronal silencing (Vyazovskiy et al. 2011) reflecting local sleep.

Importantly, only local sleep would potentially allow for independent regulation of attentional resources while a global state change would downregulate attentional resources concomitantly. Hence, the strong interaction between N2pc and HFA speaks in favor of brief periods of local sleep as single units usually do only during NREM sleep (Vyazovskiy et al. 2011; Siclari et al. 2017) even in the absence of signs of drowsiness. The HFA, a localized index of functionally selective activity (Crone et al. 1998; Miller et al. 2007) and most likely reflecting multi-unit activity vanishes during MW in regions strongly responding to stimulation. In addition, in sleep restricted humans waking EEG typically shows increased low-frequency power (slow wave activity – SWA) reflecting the duration of prior wakefulness (Finelli et al. 2000; Leemburg et al. 2010; Vyazovskiy et al. 2011) and a homologue phenomenon to silencing neurons in brain regions disproportionately used during waking (Rector et al. 2009), and involved in prior learning (Hung et al. 2013). Both strong signatures of local sleep – i.e., HFA reduction and SWA increase – did not overlap spatially but occurred locally (Bellesi et al. 2014), which points at different functions.

SWA could serve as a carrier wave that allows or drives the transfer of information between structures such as the hippocampus and neocortex and occurred over centro-parietal, sensory and motor areas regions relative to the rest of the brain in a previous study (Castelnovo et al. 2016). In line with previous results, we found an increase in centro-parietal and in occipital cortex. The parallel SWA increase between these regions argues strongly for a common plasticity dependent component to sleep regulation (Murphy et al. 2011). Importantly, these signatures of local sleep occur even in subjects which are not sleep deprived (Quercia et al. 2018) and SWA, indicating sleep need (Huber et al. 2004), varies locally in time, since subjective ON and OFF periods were reported comparably distributed across the entire experiment. Hence, we can rule out the possibility that both signatures of LS only increase with time and thus without any strong relation to MW.

Local sleep periods are of behavioral relevance since they are associated with cognitive lapses (Nir et al. 2017) marked by prolonged reaction times (Bernardi et al. 2015; Nir et al. 2017), probably due to reduced stimulus-triggered activity in visual areas causing a lower-quality perceptual representation of the target stimulus (Weissman et al. 2006). Consistent with subjects experiencing attentional lapses, we also found reaction times to be substantially longer during MW. The observed motor slowing might in part explain behavioral errors in previous studies on MW as well. MW manifests behaviorally especially in highly automated task like reading or the Sustained-Attention-to-Response-Task (SART)(Smallwood et al. 2008; Seli 2016) hence behavioral decrements in SART experiments could result from a slowing of a general control of manual responses which could hypothetically be beneficial to prevent from overhasty decisions when sensory evidence is low. The important finding is that even though low in performance, subjects with stronger N2pc perform better underscoring the behavioral relevance of upregulation of attentional resources when sensory evidence is low.

Indeed, our major finding is that during local sleep the strength of SWA and neuronal silencing predicts how attentional reallocation is modulated. Previously, MW was found to positively correlate with task-irrelevant distraction indicating that MW reveals individual susceptibility to task-irrelevant distraction including both internal and external sources (Forster and Lavie 2014). Specifically, it was suggested that MW and external distraction reflect distinct, yet correlated constructs related to working memory (Unsworth and McMillan 2014). Hence, the N2pc increase is in line with previous studies showing that target-distractor disambiguation increases with distractor load (Mazza et al. 2009) and suggesting a stronger influence of distractors under momentary attention lapses (Weissman et al. 2006). These results indicate that MW does not inflict attentional decoupling (Smallwood and Schooler 2006). Given the earlier onset of HFA compared to the N2pc, the reduction in HFA during MW (worse stimulus representation) might consequently lead to the upregulation of the N2pc (more target enhancement and/or distractor suppression needed). Since experience sampling can only be applied in a subset of trials, a trial-wise measure of MW cannot be provided. Hence, we cannot dissolve the number of trials by which neuronal silencing is ahead the N2pc upregulation.

The N2pc was originally interpreted as suppression of distractors (Luck and Hillyard 1994b), but others argued that the N2pc reflects target enhancement (Eimer 1996) and is now considered a composition of overlapping processes of both target processing (target negativity, Nt) and distractor suppression (distractor positivity, Pd) (Hickey et al. 2009; Hilimire et al. 2012; Gaspar and McDonald 2014). Since we presented the target simultaneously with a color pop-out non-target in the opposite visual field, both the target selection (Nt contralateral to the target) as well as distractor suppression (Pd contralateral to the pop-out non-target) will contribute to the amplitude of the observed N2pc waveform. Importantly, we observed an enhanced N2pc when the subjects were in a state of MW. Since our stimuli always contained both laterally presented targets and distractors, we cannot unambiguously decide as to whether the enhanced N2pc was caused by a stronger target enhancement, increased distractor suppression, or both, or whether the N2pc is rather generally suppressed in the focused state. In general, the N2pc component seems to strongly depend on stimulation parameters, showing larger activation differences between hemispheres when more than one item per visual field is presented, the task requires a complex feature discrimination (compared to a simple feature detection) and the target is in the lower visual field (Luck et al. 1997). Hence, we chose our visual search display accordingly to maximize the observed N2pc amplitudes with the target being located in the lower visual field, multiple surrounding irrelevant distractor items, and a discrimination task requiring high spatial scrutiny. Most importantly, the target was always an easily detectable color pop-out item, requiring no time-consuming search process that might have smeared out N2pc responses over time. In fact, the N2pc was elicited at the expected time range of 200 msec irrespective of mental state. That is, the initial target selection was not delayed under conditions of MW. Still, there was a substantial increase in response time (about 400msec), when subjects reported to be “OFF task” which might have reflected a delayed processing of the information provided by the N2pc, or could be caused by parallel interfering processes of MW. In fact, only when participants experienced MW (OFF task), the amplitude of the N2pc was positively correlated with performance. That is, a larger N2pc, typically associated with a stronger focusing onto the target and potentially reflecting better distractor suppression (Mazza et al. 2009; Donohue et al. 2016), might have compensated for the mind wandering.

When investigating MW, a major challenge is how to reliably capture phases of reduced focusing on the task. Frequently prompting thought probes during the course of the experiments will most likely discourage MW, hence, we chose to assess the participants mental state on only 20% of the trials. As a consequence, trial numbers are inherently limited for comparing neural responses between mental states. Furthermore, participants reported for the majority of trials (51%) to be “on task”, which might be caused by the perceptually rather demanding discrimination task, or also be influenced by participants trying to respond in a socially desirable way. Nevertheless, the markers of local sleep (SWA increase, HFA reduction) match participants self-reports with being “off the task” and might also provide future measures depending less on self-report.

Our critical conclusion is that MW is strongly linked to cortical dynamics associated with local sleep and that attentional resources needed for visual search are upregulated to circumvent restrictions caused by limited sensory evidence. Occipital HFA, which shows a strong stimulus response comparable to intracranial recordings, falls out when participants have the subjective impression of being off the task, commensurate with an increase in periods of SWA increase. Attentional decoupling as predicted for being off the task is expected to produce a decrease in the N2pc (Schad et al. 2012; Christoff et al. 2016). But reduced sensory evidence compels stronger attentional allocation to key features in the environment and hence a stronger target-distractor disambiguation during MW. Hence these results indicate that MW does not lead to a global blackout of HFA but cortical regions generating the target-distractor disambiguation also flexibly reacts to internal distractions. These functional explanations indicate that expected input to visual stimulation is tracked and stronger reallocation of spatial attention is generated when sensory evidence is scarce, presumably by frontal cortical areas. In sum, we provide evidence that MW is strongly related to local sleep and establish a direct link between boosted attentional resources due to local sleep during waking.

## Acknowledgments

This work was funded by ‘Schwerpunkt Neuroforschung. MW-21 LMSP 9-2012’, funded by Land Sachsen-Anhalt. C.R. was funded by the Federal Ministry of Education and Research, Germany, grant number 13GW0095D.

## References

Andrillon T, Windt J, Silk T, Drummond SPA, Bellgrove MA, Tsuchiya N. 2019. Does the Mind Wander When the Brain Takes a Break? Local Sleep in Wakefulness, Attentional Lapses and Mind-Wandering. Front Neurosci. 13:1–10.

Bartoli E, Bosking W, Li Y, Beauchamp MS, Yoshor D, Foster B. 2019. Distinct Narrow and Broadband Gamma Responses in Human Visual Cortex. bioRxiv. 572313.

Bellesi M, Riedner BA, Garcia-Molina GN, Cirelli C, Tononi G. 2014. Enhancement of sleep slow waves: Underlying mechanisms and practical consequences. Front Syst Neurosci. 8:1–17.

Bernardi G, Siclari F, Yu I, Zennig C, Bellesi M, Ricciardi E, Cirelli C, Ghilardi MF, Pietrini P, Tononi G. 2015. Neural and behavioral correlates of extended training during sleep deprivation in humans: Evidence for local, task-specific effects. J Neurosci. 35:4487–4500.

Boehler CN, Tsotsos JK, Schoenfeld MA, Heinze H-J, Hopf J-M. 2011. Neural Mechanisms of Surround Attenuation and Distractor Competition in Visual Search. J Neurosci. 31:5213–5224.

Brainard DH. 1997. The Psychophysics Toolbox. Spat Vis. 10:433–436.

Brandmeyer T, Delorme A. 2018. Reduced mind wandering in experienced meditators and associated EEG correlates. Exp Brain Res. 236:2519–2528.

Burke JF, Long NM, Zaghloul KA, Sharan AD, Sperling MR, Kahana MJ. 2014. Human intracranial high-frequency activity maps episodic memory formation in space and time. Neuroimage. 85:834–843.

Carriere JSA, Cheyne JA, Smilek D. 2008. Everyday attention lapses and memory failures: The affective consequences of mindlessness. Conscious Cogn. 17:835–847.

Castelnovo A, Riedner BA, Smith RF, Tononi G, Boly M, Benca RM. 2016. Scalp and Source Power Topography in Sleepwalking and Sleep Terrors: A High-Density EEG Study. Sleep. 39:1815–1825.

Christoff K, Irving ZC, Fox KCR, Spreng RN, Andrews-Hanna JR. 2016. Mind-wandering as spontaneous thought: a dynamic framework. Nat Rev Neurosci. 17:718–731.

Coon WG, Schalk G. 2016. A method to establish the spatiotemporal evolution of task-related cortical activity from electrocorticographic signals in single trials. J Neurosci Methods. 271:76–85.

Crone N, Miglioretti DL, Gordon B, Lesser RP. 1998. Functional mapping of human sensorimotor cortex with electrocorticographic spectral analysis. II. Event-related synchronization in the gamma band. Brain. 121:2301–2315.

Donohue SE, Hopf J-M, Bartsch M V., Schoenfeld MA, Heinze H-J, Woldorff MG. 2016. The Rapid Capture of Attention by Rewarded Objects. J Cogn Neurosci. 28:529–541.

Dürschmid S, Edwards E, Reichert C, Dewar C, Hinrichs H, Heinze HJ, Kirsch HE, Dalal SS, Deouell LY, Knight RT. 2016. Hierarchy of prediction errors for auditory events in human temporal and frontal cortex. Proc Natl Acad Sci U S A. 113:6755–6760.

Eimer M. 1996. The N2pc component as an indicator of attentional selectivity. Electroencephalogr Clin Neurophysiol. 99:225–234.

Finelli LA, Baumann H, Borbély AA, Achermann P. 2000. Dual electroencephalogram markers of human sleep homeostasis: Correlation between theta activity in waking and slow-wave activity in sleep. Neuroscience. 101:523–529.

Forster S, Lavie N. 2014. Distracted by your mind? Individual differences in distractibility predict mind wandering. J Exp Psychol Learn Mem Cogn. 40:251–260.

Gaspar JM, McDonald JJ. 2014. Suppression of salient objects prevents distraction in visual search. J Neurosci. 34:5658–5666.

Gerber EM, Golan T, Knight RT, Deouell LY. 2017. Cortical representation of persistent visual stimuli. Neuroimage. 161:67–79.

Golan T, Davidesco I, Meshulam M, Groppe DM, Mégevand P, Yeagle EM, Goldfinger MS, Harel M, Melloni L, Schroeder CE, Deouell LY, Mehta AD, Malach R. 2016. Human intracranial recordings link suppressed transients rather than “filling-in” to perceptual continuity across blinks. Elife. 5:1–28.

Golan T, Davidesco I, Meshulam M, Groppe DM, Mégevand P, Yeagle EM, Goldfinger MS, Harel M, Melloni L, Schroeder CE, Deouell LY, Mehta AD, Malach R. 2017. Increasing suppression of saccade-related transients along the human visual hierarchy. Elife. 6:1–15.

He J, Becic E, Lee YC, McCarley JS. 2011. Mind wandering behind the wheel: Performance and oculomotor correlates. Hum Factors. 53:13–21.

Helfrich RF, Fiebelkorn IC, Szczepanski SM, Lin JJ, Parvizi J, Knight RT, Kastner S. 2018. Neural Mechanisms of Sustained Attention Are Rhythmic. Neuron. 99:854-865.e5.

Hickey C, Di Lollo V, McDonald JJ. 2009. Electrophysiological indices of target and distractor processing in visual search. J Cogn Neurosci. 21:760–775.

Hilimire MR, Hickey C, Corballis PM. 2012. Target resolution in visual search involves the direct suppression of distractors: Evidence from electrophysiology. Psychophysiology. 49:504–509.

Hilimire MR, Mounts JRW, Parks NA, Corballis PM. 2011. Dynamics of target and distractor processing in visual search: Evidence from event-related brain potentials. Neurosci Lett. 495:196–200.

Hopf J-M, Luck SJ, Girelli M, Hagner T, Mangun GR, Scheich H, Heinze H-J. 2000. Neural Sources of Focused Attention in Visual Search. Cereb Cortex. 10:1233–1241.

Huber R, Felice Ghilardi M, Massimini M, Tononi G. 2004. Local sleep and learning. Nature. 430:78–81.

Hung C-S, Sarasso S, Ferrarelli F, Riedner B, Ghilardi MF, Cirelli C, Tononi G. 2013. Local Experience-Dependent Changes in the Wake EEG after Prolonged Wakefulness. Sleep. 36:59–72.

Kam JWY, Dao E, Farley J, Fitzpatrick K, Smallwood J, Schooler JW, Handy TC. 2011. Slow fluctuations in attentional control of sensory cortex. J Cogn Neurosci. 23:460–470.

Kam JWY, Solbakk AK, Endestad T, Meling TR, Knight RT. 2018. Lateral prefrontal cortex lesion impairs regulation of internally and externally directed attention. Neuroimage. 175:91–99.

Kleiner M, Brainard D, Pelli D, Ingling A, Murray R, Broussard C. 2007. What’s new in Psychtoolbox-3 ?

Kucyi A, Salomons T V., Davis KD. 2013. Mind wandering away from pain dynamically engages antinociceptive and default mode brain networks. Proc Natl Acad Sci. 110:18692–18697.

Kupers E, Wang H, Amano K, Kay K, Heeger D, Winawer J. 2017. A non-invasive, quantitative study of broadband spectral responses in human visual cortex. Broadband Spectr responses Vis cortex Reveal by a new MEG denoising algorithm. 108993.

Lee B, Martin P, Valberg A. 1988. The physiological basis of heterochromatic flicker photometry. J Physiol. 323–347.

Leemburg S, Vyazovskiy V V., Olcese U, Bassetti CL, Tononi G, Cirelli C. 2010. Sleep homeostasis in the rat is preserved during chronic sleep restriction. Proc Natl Acad Sci U S A. 107:15939–15944.

Leszczynski M, Chaieb L, Reber TP, Derner M, Axmacher N, Fell J. 2017. Mind wandering simultaneously prolongs reactions and promotes creative incubation. Sci Rep. 7:1–9.

Liu J, Newsome WT. 2006. Local field potential in cortical area MT: Stimulus tuning and behavioral correlations. J Neurosci. 26:7779–7790.

Luck SJ. 2005. An Introduction to the Event-Related Potential Technique. Cambridge, MA: MIT Press.

Luck SJ, Girelli M, McDermott MT, Ford MA. 1997. Bridging the Gap between Monkey Neurophysiology and Human Perception: An Ambiguity Resolution Theory of Visual Selective Attention, Cognitive Psychology.

Luck SJ, Hillyard SA. 1994a. Electrophysiological correlates of feature analysis during visual search. Psychophysiology. 31:291–308.

Luck SJ, Hillyard SA. 1994b. Spatial Filtering During Visual Search: Evidence From Human Electrophysiology. J Exp Psychol Hum Percept Perform. 20:1000–1014.

Manning JR, Jacobs J, Fried I, Kahana MJ. 2009. Broadband Shifts in Local Field Potential Power Spectra Are Correlated with Single-Neuron Spiking in Humans. 29:13613–13620.

Mazza V, Turatto M, Caramazza A. 2009. Attention selection, distractor suppression and N2pc. CORTEX. 45:879–890.

Miller KJ, denNijs M, Shenoy P, Miller JW, Rao RPN, Ojemann JG. 2007. Real-time functional brain mapping using electrocorticography. Neuroimage. 37:504–507.

Miller KJ, Honey CJ, Hermes D, Rao RPN, DenNijs M, Ojemann JG. 2014. Broadband changes in the cortical surface potential track activation of functionally diverse neuronal populations. Neuroimage. 85:711–720.

Miller KJ, Sorensen LB, Ojemann JG, Den Nijs M. 2009. Power-law scaling in the brain surface electric potential. PLoS Comput Biol. 5.

Mukamel R, Gelbard H, Arieli A, Hasson U, Fried I, Malach R. 2005. Neuroscience: Coupling between neuronal firing, field potentials, and fMRI in human auditory cortex. Science (80-). 309:951–954.

Murphy M, Huber R, Esser S, A. Riedner B, Massimini M, Ferrarelli F, Felice Ghilardi M, Tononi G. 2011. The Cortical Topography of Local Sleep. Curr Top Med Chem. 11:2438–2446.

Müsch K, Hamamé CM, Perrone-Bertolotti M, Minotti L, Kahane P, Engel AK, Lachaux JP, Schneider TR. 2014. Selective attention modulates high-frequency activity in the face-processing network. Cortex. 60:34–51.

Nir Y, Andrillon T, Marmelshtein A, Suthana N, Cirelli C, Tononi G, Fried I. 2017. Selective neuronal lapses precede human cognitive lapses following sleep deprivation. Nat Med. 23:1474–1480.

Pelli DG. 1997. The VideoToolbox software for visual psychophysics: transforming numbers into movies. Spat Vis. 10:437–442.

Perrone-Bertolotti M, El Bouzaïdi Tiali S, Vidal JR, Petton M, Croize AC, Deman P, Rheims S, Minotti L, Bhattacharjee M, Baciu M, Kahane P, Lachaux JP. 2020. A real-time marker of object-based attention in the human brain. A possible component of a “gate-keeping mechanism” performing late attentional selection in the Ventro-Lateral Prefrontal Cortex. Neuroimage. 210:116574.

Privman E, Malach R, Yeshurun Y. 2013. Modeling the electrical field created by mass neural activity. Neural Networks. 40:44–51.

Quercia A, Zappasodi F, Committeri G, Ferrara M. 2018. Local use-dependent sleep in wakefulness links performance errors to learning. Front Hum Neurosci. 12:1–17.

Ray S, Maunsell JHR. 2011. Different Origins of Gamma Rhythm and High-Gamma Activity in Macaque Visual Cortex. PLoS Biol. 9:e1000610.

Rector DM, Schei JL, Van Dongen HPA, Belenky G, Krueger JM. 2009. Physiological markers of local sleep. Eur J Neurosci. 29:1771–1778.

Schad DJ, Nuthmann A, Engbert R. 2012. Your mind wanders weakly, your mind wanders deeply: Objective measures reveal mindless reading at different levels. Cognition. 125:179–194.

Seli P. 2016. The Attention-Lapse and Motor Decoupling accounts of SART performance are not mutually exclusive. Conscious Cogn. 41:189–198.

Siclari F, Baird B, Perogamvros L, Bernardi G, LaRocque JJ, Riedner B, Boly M, Postle BR, Tononi G. 2017. The neural correlates of dreaming. Nat Neurosci. 20:872–878.

Smallwood J, Beach E, Schooler JW, Handy TC. 2008. Going AWOL in the Brain: Mind Wandering Reduces Cortical Analysis of External Events. J Cogn Neurosci. 20:458–469.

Smallwood J, Schooler JW. 2006. The restless mind. Psychol Bull. 132:946–958.

Szczepanski SM, Crone NE, Kuperman RA, Auguste KI, Parvizi J, Knight RT. 2014. Dynamic Changes in Phase-Amplitude Coupling Facilitate Spatial Attention Control in Fronto-Parietal Cortex. PLoS Biol. 12:e1001936.

Szczepanski SM, Kastner S. 2013. Shifting Attentional Priorities: Control of Spatial Attention through Hemispheric Competition. J Neurosci. 33:5411–5421.

Szczepanski SM, Knight RT. 2014. Insights into Human Behavior from Lesions to the Prefrontal Cortex. Neuron. 83:1002–1018.

Unsworth N, McMillan BD. 2014. Similarities and differences between mind-wandering and external distraction: A latent variable analysis of lapses of attention and their relation to cognitive abilities. Acta Psychol (Amst). 150:14–25.

Vyazovskiy V V., Harris KD. 2013. Sleep and the single neuron: The role of global slow oscillations in individual cell rest. Nat Rev Neurosci. 14:443–451.

Vyazovskiy V V, Olcese U, Hanlon EC, Nir Y, Cirelli C, Tononi G. 2011. Local sleep in awake rats. Nature. 472:443–447.

Weissman DH, Roberts KC, Visscher KM, Woldorff MG. 2006. The neural bases of momentary lapses in attention. Nat Neurosci. 9:971–978.

Westner BU, Dalal SS, Hanslmayr S, Staudigl T. 2018. Across-subjects classification of stimulus modality from human MEG high frequency activity. PLoS Comput Biol. 14:1–14.

Yanko MR, Spalek TM. 2014. Driving with the wandering mind: The effect that mind-wandering has on driving performance. Hum Factors. 56:260–269.

